# Soils from different landscape elements diverge in response to multiple global change factors

**DOI:** 10.64898/2026.09.01.748383

**Authors:** Tian Hu, Mohan Bi, Matthias C. Rillig

## Abstract

Global change factors (GCFs) are known to affect terrestrial ecosystems across a range of land-use types, including farmlands, grasslands, and forests. However, it remains unclear whether different landscape elements within the same region respond differently to the same global change pressures. Here, we investigated this question using four soils collected from co-located farmland, grassland, pine forest, and oak forest, representing distinct landscape elements under the same regional climatic conditions. Each soil was exposed to one of six individual GCFs, warming, drought, nitrogen deposition, salinity, microplastics, and antibiotics, as well as to all six factors combined. We found that landscape elements exhibited strongly divergent responses to the same GCFs. The effects on soil functions also varied among soils under combined stress, with responses diverging from different null-model predictions depending on soils and response variable. Moreover, landscape-element specific response patterns became more pronounced under multiple concurrent stressors. Overall, our findings show that landscape heterogeneity represents a mosaic of different capacities to resist and respond to global change, even under shared climatic and geographic conditions. Global change assessments and ecosystem models should therefore better account for landscape-level heterogeneity, and management strategies aimed at enhancing ecosystem resilience should be tailored to individual landscape elements.

## 1. Introduction

Global change factors (GCFs), including warming, nutrient deposition, salinity, or emerging pollutants, are increasingly affecting terrestrial ecosystems worldwide^1^. These environmental stressors can substantially change soil physicochemical properties, microbial activity, and ecosystem functioning^2^. While numerous experimental studies have investigated the effects of global change factors on individual soil ecosystems, natural environments are rarely exposed to stressors in isolation. Ecosystems are interconnected systems linked through material and organism transport, and contain multiple interacting biological and physicochemical processes that can buffer simultaneous environmental stressors^3,4^. Comparative studies across landscapes and ecosystems are necessary to understand the impacts of global change.

Previous studies have shown that global change factors can significantly affect soil ecosystems across different landscape types, including forests, grasslands, and agricultural ecosystems^5–13^. In addition, several meta-analyses have demonstrated that the impacts of global change factors often differ among ecosystem types due to variation in vegetation cover, land-use history, and soil physicochemical properties^14–16^. Soil microbial communities also respond differently to global change across ecosystem types^15^. For example, the sensitivity of nitrogen loss processes to individual global change factor varies substantially among forests, grasslands, and agricultural systems, and is largely regulated by local climate and soil conditions^14^. Moreover, high-urbanity and low-urbanity soils were found to react differently to multiple stressors^17^. These findings suggest that soil responses to environmental stress could be context dependent and regulated by intrinsic ecosystem characteristics. Contaminant risks and ecosystem responses are landscape-element dependent, because patches differ in their capacity to capture, transform, and buffer environmental stressors^18^. Therefore, it is important to examine the stressor effects across landscape elements and ecological scales^19^. Despite previous work showed that different ecosystems may respond differently to global change, most existing studies either focus on individual ecosystem types or synthesize independent studies through meta-analysis^20^. To our knowledge, no controlled experimental study has directly compared the responses of soils from contrasting landscape elements within the same geographic and climatic context to an identical set of global change factors. As a result, it remains unclear to what extent landscape context regulate ecosystem sensitivity and functional responses to multiple environmental stressors.

To address this knowledge gap, we conducted a controlled microcosm experiment using soils collected from four directly adjacent landscape elements, including agricultural land, grassland, pine forest, and oak forest. Six global change factors warming, drought, nitrogen deposition, salinity, microplastics, antibiotics were applied individually and in combination. We measured multiple soil physicochemical properties, including soil pH, electrical conductivity (EC), water-stable aggregates (WSA), and soil biological and functional responses including litter decomposition rate, and soil respiration. We aimed to determine how soil from different landscape elements react differently to single and multiple global change factors. We hypothesized that (1) soils from different landscape elements will exhibit divergent responses to global change factors, with soil origin shaping the magnitude and direction of ecosystem sensitivity; (2) jointly acting global change factors will generate non-additive effects that deviate from single-factor null model predictions, with interaction patterns varying among landscape elements; and (3) variation in soil response patterns among landscape elements is amplified under multiple stressors.

## 2. Methods

### 2.1. Experimental design

Soils were collected in July 2025 from the Forschungsstation Linde, a research site managed by the Zwillenberg–Tietz Foundation, located in the federal State of Brandenburg near Berlin (**Figure S1**). Site 1 is an organically-managed farmland field, which was planted with winter rye at the time of sampling. The field had previously been sown with summer rye in spring 2024 and field mustard as a catch crop in autumn 2023. The soil is a Bändersand-Rumpfrosterde. Site 2 was located in a 81 to 100 years old oak forest, and the soil is Kersdorfer Sand-Ranker. Site 3 was situated in an over 100 year-old Scots pine forest (*Pinus sylvestris*), and the soil is Neudarßer Sand-Saumpodsol. Site 4 is a grassland field, which had been cultivated with alfalfa for approximately four years prior to sampling. The grassland was mown two to three times annually, and the harvested biomass was used as silage for dairy cattle. The soil is Schwarzheider Tieflehm-Fahlerde. These four sites represent distinct landscape elements, with the largest difference between two sites only around 2 km. Detailed physicochemical properties of each soil are provided in **Table S1**^21^.

Each soil had a control treatment (10 replicates), and was subjected to six single-factor treatments in which each global change factor was applied individually (8 replicates per treatment), and one six-factor treatment in which all global change factors were applied simultaneously (10 replicates). Six global change factors were selected to represent a broad spectrum of environmental stressors: warming, drought, nitrogen deposition, salinity, microplastics, and antibiotics. These factors cover climatic changes (temperature and moisture), inorganic and organic pollutants, particulate pollutants, and emerging contaminants. In total, each soil comprised 68 experimental units, resulting in 272 microcosms across all four soils (**Figure 1**).

**Fig. 1.**
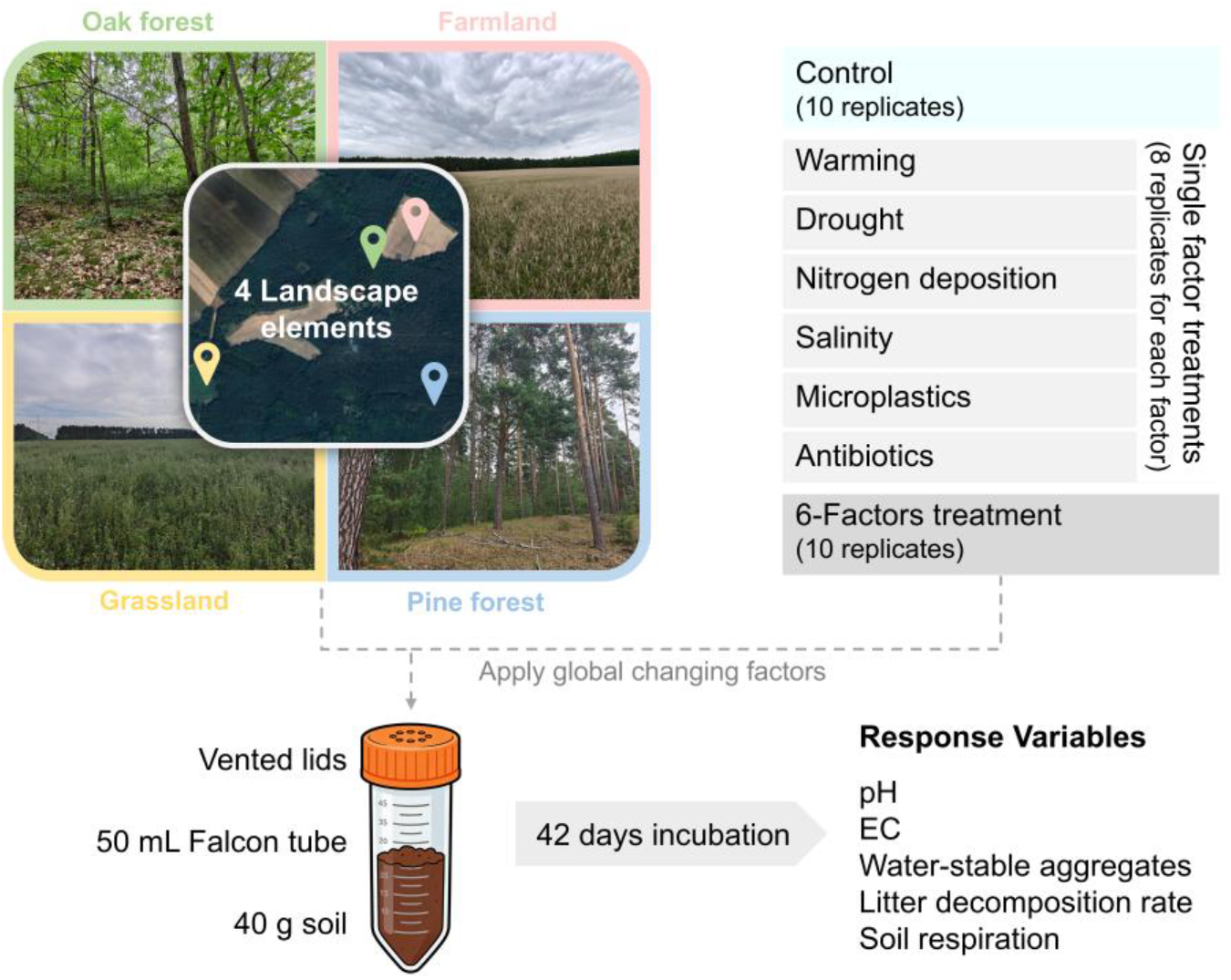
Experimental design. Four landscape elements were included: oak forest, farmland, grassland, and pine forest. Six global change factor (GCF) treatments—warming, drought, nitrogen deposition, salinity, microplastics, and antibiotics—were applied individually with eight replicates each, together with a combined six-factor treatment with 10 replicates and an untreated control with 10 replicates. For each experimental unit, 40 g of soil was incubated in a 50-mL Falcon tube with a vented lid for 42 days. After incubation, soil samples were harvested and analyzed for pH, electrical conductivity, water-stable aggregates, litter decomposition rate, and soil respiration. Photographs were taken on the day of sampling.

### 2.2. Soil preparation and microcosm setup

For each landscape element, topsoil (0–10 cm depth) was sampled after removing surface litter. All soil samples were air-dried and sieved through a 2 mm mesh to remove visible roots and stones.

Microcosms were established in 50 mL conical mini-bioreactors (Product Nr: 431720, Corning®, USA) with a vented film, which allows gas exchange but prevents microbial contamination. Each contained 40 g dry weight (dw) of soil. A sterilized tea bag was vertically inserted into the center of each microcosm to assess litter decomposition. Soil moisture was adjusted to 60% water holding capacity (WHC) (or 30% WHC for drought treatments) using distilled water.

### 2.3. Application of GCFs and harvest

The six global change factors were applied at environmentally relevant levels commonly used in previous experimental studies^22,23^. Treatments included warming (+5 °C above ambient temperature), drought (30% WHC versus 60% WHC in controls), nitrogen deposition (100 kg N ha⁻¹ yr⁻¹ with NH₄NO₃), salinity (4.0 dS m⁻¹ with NaCl), microplastic contamination (0.1% w/w polyethylene particles), and antibiotic pollution (3 mg oxytetracycline kg⁻¹ soil). Warming was achieved using heating cables (PT2011, Exo Terra, Germany) regulated by individual temperature controllers (ETC-902, VOLTCRAFT, Germany) for each experimental unit.

To achieve the homogeneous application of GCF treatments, we used a “loading soil”, a small amount (5.0 g) of soil sterilized at 121 °C for 20 minutes. This sterilized fraction was used as “loading soil” to mixed chemical-based GCF treatments evenly into the experimental soil; the sterilization of this small amount of soil was done to avoid strong effect on soil microbiota^1^. For soluble stressors nitrogen deposition and antibiotic pollution, NH₄NO₃ (≥98%, p.A., ACS. Roth GmbH, Karlsruhe, Germany, article K299.1) and oxytetracycline (Sigma-Aldrich, MO, USA, catalog #PHR1537) solutions were prepared in distilled water, and 100 μL of solution was added to the loading soil. Polyethylene particles (mean particle size 150 μm, Thermo Scientific, USA, catalog #043951.18) and NaCl (≥99%,, Ph. Eur., USP. Roth GmbH, Karlsruhe, Germany. article P029.2) were added directly for the microplastic and salinity treatments, respectively. Soil mixtures were then homogenized by shaking with a shaking machine (Product Nr: 541-21009-00, Reax2, Heidolph Instrument GmbH & Co. KG, Schwabach, Germany) at 80 rpm for 30 min before being transferred to the microcosm bioreactors.

Bioreactors were placed in sand-filled cups to minimize heat exchange among experimental units and placed in a fully randomized fashion inside a climate-controlled chamber. The incubation was conducted in the dark at 20°C and 60% relative humidity. Weight of mini-bioreactors was measured individually each week, and distilled water was added to restore the original weight and maintain a constant water content throughout the experiment. After 42 days of incubation, all microcosms were destructively sampled. Tea bags were carefully removed, oven-dried at 60 °C, and weighed to determine litter decomposition rates. The remaining soil was homogenized and air-dried at room temperature for physicochemical analyses.

Soil responses were assessed in terms of physicochemical properties and biological function. Physicochemical measurements included soil pH, electrical conductivity, and water-stable aggregates. Biological and functional responses included litter decomposition rate and soil respiration. For soil pH, 5.0 g of air-dried soil was thoroughly mixed with 25 mL of 0.01 M CaCl₂ solution in a 50-mL centrifuge tube using a vortex mixer. The suspension was then centrifuged at 4600 rpm for 10 min at room temperature, and pH was measured using a pH meter (Hanna Instruments, Smithfield, USA). For soil EC, 5.0 g of air-dried soil was mixed with 25 mL of distilled water in a 50-mL centrifuge tube and shaken at 250 rpm for 1 h and then measured (SevenEasy pH-Meter Mettler-Toledo). The percentage of water-stable aggregates was determined using a wet-sieving method^24^. Briefly, 4.0 g of soil was placed on a 0.25-mm mesh sieve, rewetted by capillary action with deionized water, and subsequently subjected to wet sieving using a sieving machine (Agrisearch Equipment, Eijkelkamp, Giesbeek, Netherlands). The percentage of water-stable aggregates was calculated as: %WSA = (water stable fraction-coarse matter)/(4.0 g-coarse matter). Soil respiration was measured as the change in CO₂ concentration (ppm)^25^. Before each measurement, the headspace of each tube was flushed with CO₂-free air for 5 min to standardize the initial conditions. A 1-mL air sample was then collected from the headspace and injected into an infrared gas analyzer (Li-COR 6400XT). After 2 h of incubation, a second 1-mL air sample was collected. Soil respiration was calculated as the difference between the final and initial CO₂ concentrations. Litter decomposition rate was determined using the tea bag index method^26^. Briefly, a sealed tea bag with a mesh size of 38 μm containing 300.0 mg-dw of tea biomass was placed in the soil. Litter decomposition rate was calculated from the proportional loss of tea biomass based on its dry weight before and after incubation.

### 2.4. Statistical analysis

All statistical analyses were conducted in R (version 4.5.2). To examine variation in soil responses to GCF treatments across soils, we fitted a general linear model (GLM) for each soil response, with soil, treatment, and their interaction included as explanatory terms. To account for baseline differences among soils, treatment effects were expressed relative to the corresponding soil-type-specific control mean, calculated as the observed response minus the control mean. The contribution of soil, treatment, and their interaction to variation in each response were evaluated using Type 2 sums of squares, and the statistical significance of these model terms was assessed using ANOVA from the car R package. To compare each treatment with the corresponding control, Dunnett’ s multiple comparison tests were performed on the raw data using the emmeans package.

Treatment effects were further quantified using estimation statistics implemented in the dabestr package. For each soil and response variable, each treatment was compared with the corresponding control to calculate the effect size with 95% confidence intervals based on 5,000 bootstrap resamples. Standardized effect sizes were subsequently calculated as,

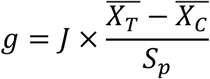

where g is Hedges’ g effect size, J is small-sample bias correction factor, *X̂_T_* is treatment group mean, *X̂_c_* is control group mean, and *S_p_* is pooled standard deviation. For each Hedges’ g estimate, 95% confidence intervals were obtained using the same bootstrap resampling.

To evaluate whether the responses to combined global change factors could be explained by the effects of individual stressors, null model analysis was performed for each soil and response variable. Three null hypotheses (additive, multiplicative, and dominative effects) were generated from the six single-factor treatments and compared with the observed six-factor treatment. For the additive model,

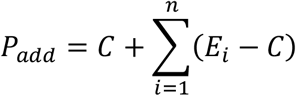

was calculated, where *P_add_* is the predicted response under the additive model, *C* is the control group mean, *E_i_* is the mean response of the *i*th individual GCF treatment, and *n* is the number of individual GCFs included in the combined treatment. For multiplicative model,

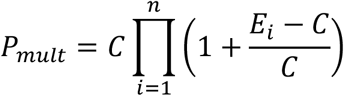

was calculated. For the dominative model,

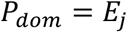

was calculated, where *P_dom_* is the predicted response under the dominative model, *E_j_* is the mean response of the individual GCF producing the largest absolute effect, and represent the GCF with the largest absolute deviation from the control.

To evaluate overall response patterns among soils, we conducted principal component analysis (PCA) and non-metric multidimensional scaling (NMDS) using the Hedges’ g effect size matrix. PCA was conducted using the FactoMineR, while NMDS was performed using the vegan package based on Euclidean distances. Permutational multivariate analysis of variance (PERMANOVA) was subsequently conducted using the adonis2() function in vegan to test the effects of soil, treatment, and their interaction on multivariate effect-size patterns, with significance assessed using 999 permutations.

Pearson correlation coefficients among the 19 soil physicochemical properties were calculated for each of the four soils. To assess the relationships between soil physicochemical properties and the treatment-induced effect sizes of the five response variables, Mantel test was performed using the linkET package. For each response variable, effect sizes were calculated as the mean Hedges’ g across the seven experimental treatments within each soil. Mantel tests were conducted separately between each response variable and each of the 19 soil properties, based on Euclidean distance matrices, using Pearson’s product-moment correlation with 999 permutations (as implemented via the vegan package within linkET). Mantel’s r and p-values were presented and visualized alongside the soil property correlation matrix using the linkET package.

## 3. Results and discussion

### 3.1. Landscape elements have different responses to global change factors

General linear models revealed significant soil × GCF treatment interactions for all five response variables (*p* < 0.001), indicating that effects of the same global change factors varied significantly among soils from the different landscape elements (**Table S2**).

Responses to individual GCFs were strongly soil-dependent. Most GCF treatments reduced water-stable aggregates across the four soils (**Figure 2**). Environmental stress can weaken aggregate stability by disrupting soil structure and reducing microbial production of extracellular polymers and fungal hyphae that bind soil particles together^27,28^. Nitrogen deposition significantly increased soil respiration in farmland and oak forest soils, while drought significantly increased respiration in grassland soil (**Figure 3**). Nitrogen deposition stimulated respiration in some soils, likely because low nitrogen inputs can alleviate nutrient limitation and enhance microbial activity^29^. Other treatments significantly reduced soil respiration. Drought or soil drying caused by warming usually inhibits soil respiration because of suppressed microbial activity^30^. Salinity and antibiotics reduced respiration due to their osmotic and toxic effects on soil microorganisms^31,32^. Microplastics can have a negative impact by potentially disrupting nutrient transportation or affecting soil bacterial communities^33^. Almost all GCF treatments reduced decomposition, and none significantly increased it (**Figure 4**). This suggests that decomposition occurs under specific environmental conditions, making it more susceptible to disruption than enhancement^34^. The strongest single-factor suppression was caused by salinity in farmland and grassland, but drought in oak and pine forest soils. Microbial communities in farmland and grassland soils are less tolerant to osmotic stress^31^, while forest decomposition relies more strongly on stable moisture conditions^35^.

**Fig. 2.**
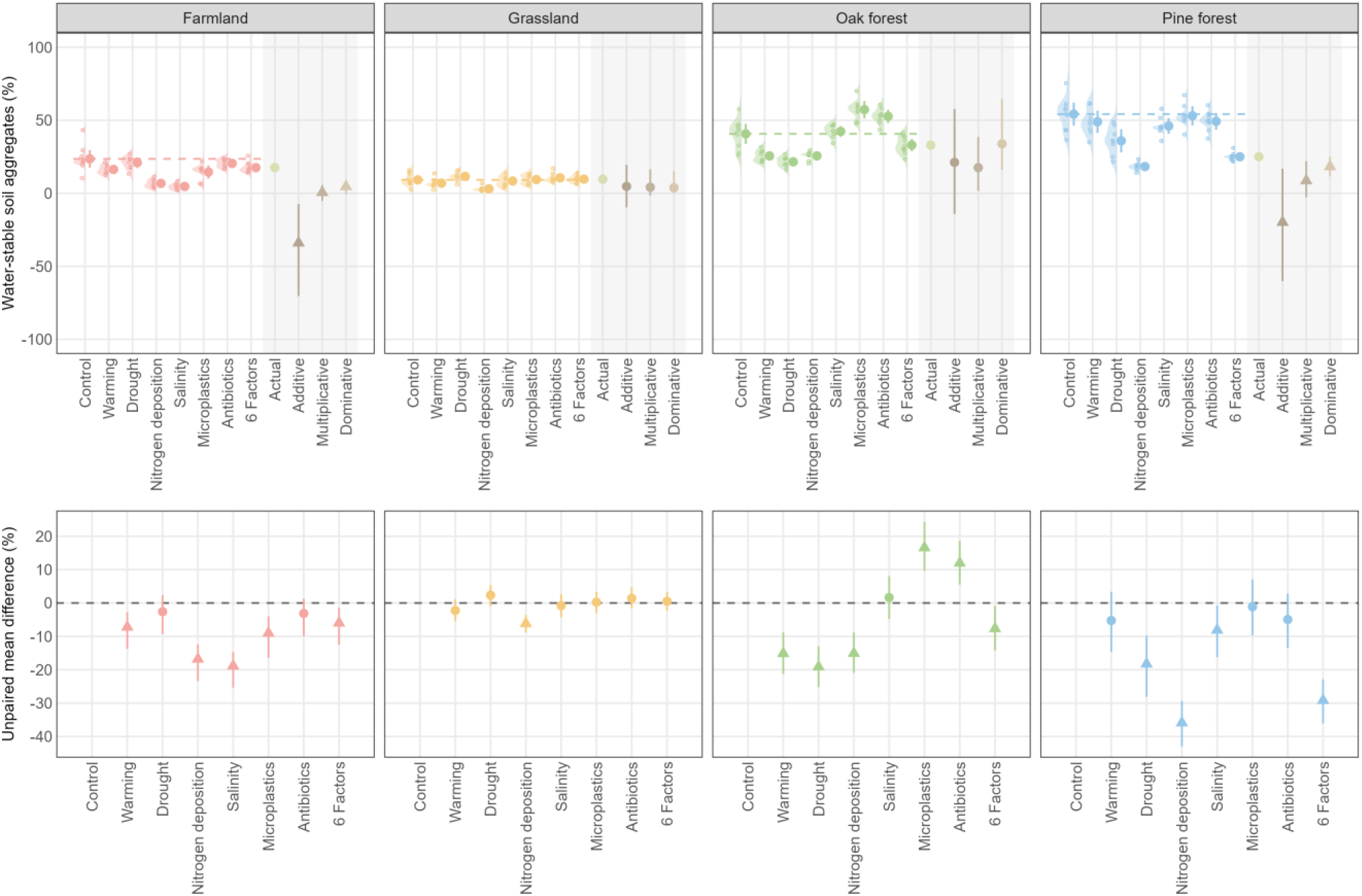
Responses of water-stable aggregates to individual and combined global change factor treatments across four landscape elements. Top panels show the raw data distributions (ridge plots and jittered points) together with treatment means ± 95% confidence intervals. The dashed horizontal line indicates the mean water-stable aggregates value of the control group. The grey-shaded area shows the observed (“Actual”) water-stable aggregates value under the combined six-factor treatment and the values predicted by three null models (Additive, Multiplicative, and Dominative); triangles indicate a significant difference between the null model prediction and the actual value, while circles indicate no significant difference. Bottom panels show the unpaired mean difference in water-stable aggregates between each treatment and the control group, with circles/triangles representing the effect size (mean difference) and its 95% confidence interval; triangles indicate a significant difference from the control (95% CI excluding zero), while circles indicate a non-significant difference.

**Fig. 3.**
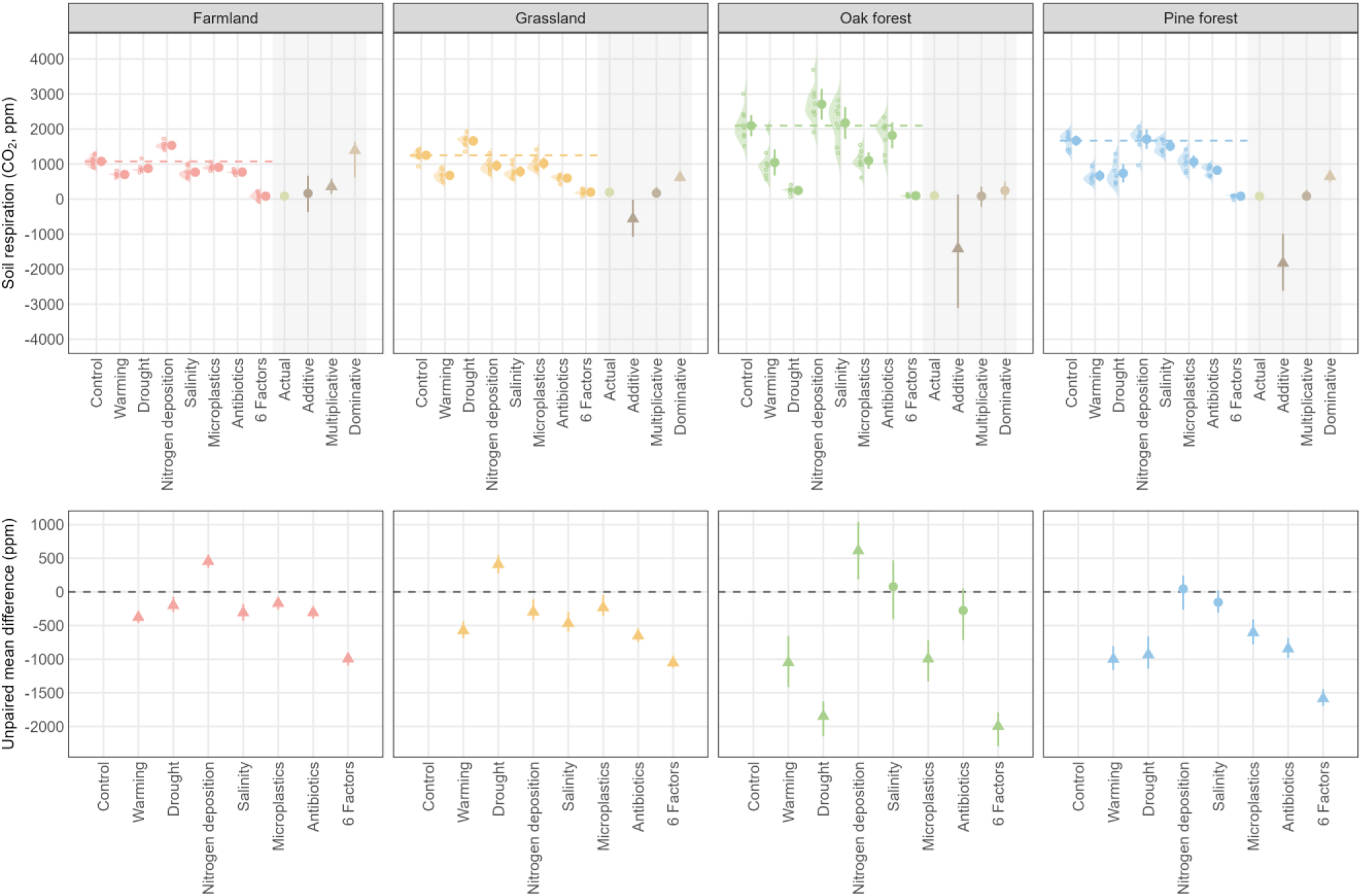
Responses of soil respiration rate to individual and combined global change factor treatments across four landscape elements. Top panels show the raw data distributions (ridge plots and jittered points) together with treatment means ± 95% confidence intervals. The dashed horizontal line indicates the mean soil respiration rate of the control group. The grey-shaded area shows the observed (“Actual”) soil respiration rate under the combined six-factor treatment and the values predicted by three null models (Additive, Multiplicative, and Dominative); triangles indicate a significant difference between the null model prediction and the actual value, while circles indicate no significant difference. Bottom panels show the unpaired mean difference in soil respiration rate between each treatment and the control group, with circles/triangles representing the effect size (mean difference) and its 95% confidence interval; triangles indicate a significant difference from the control (95% CI excluding zero), while circles indicate a non-significant difference.

**Fig. 4.**
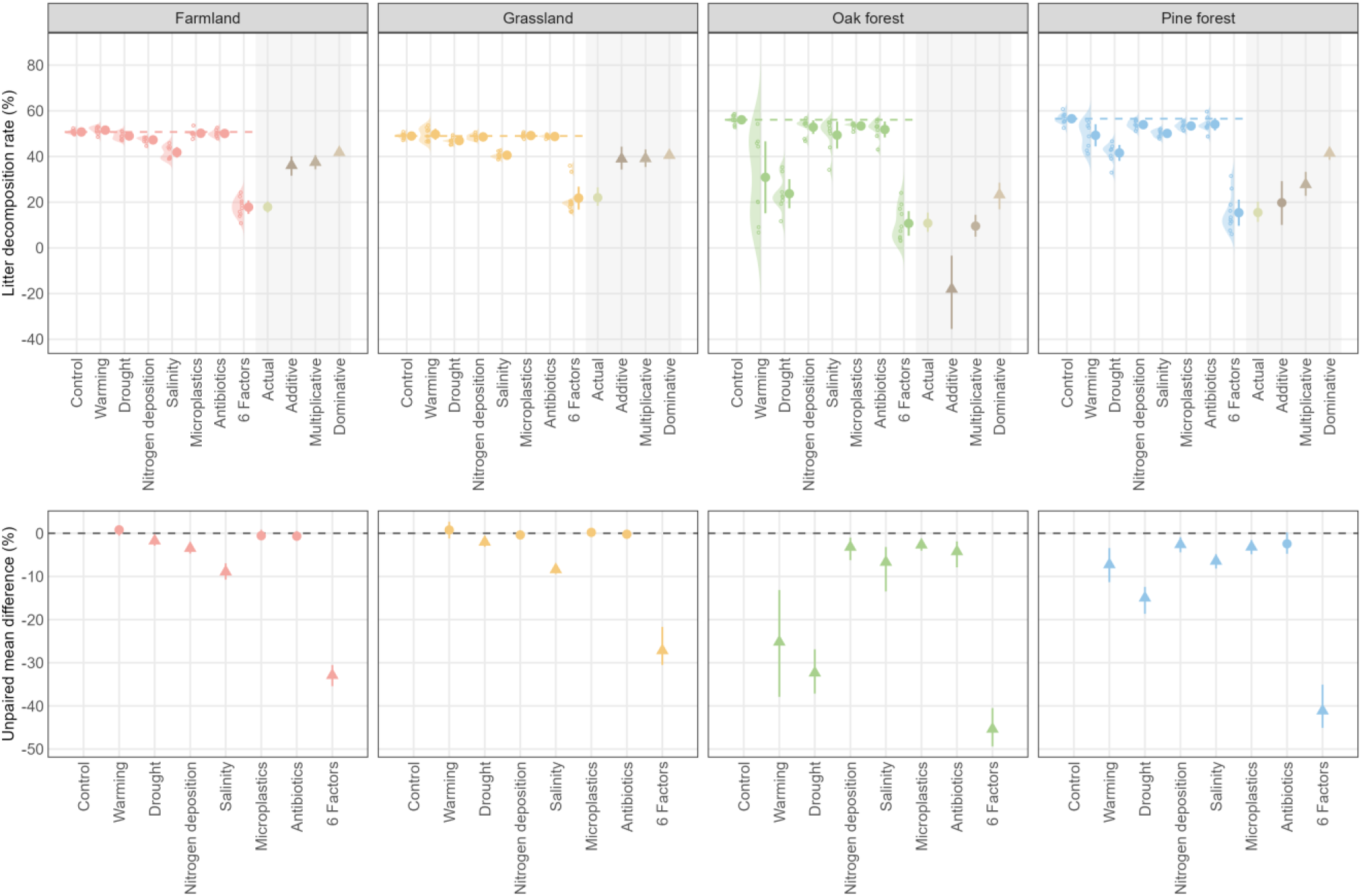
Responses of litter decomposition rate to individual and combined global change factor treatments across four landscape elements. Top panels show the raw data distributions (ridge plots and jittered points) together with treatment means ± 95% confidence intervals. The dashed horizontal line indicates the mean litter decomposition rate of the control group. The grey-shaded area shows the observed (“Actual”) litter decomposition rate under the combined six-factor treatment and the values predicted by three null models (Additive, Multiplicative, and Dominative); triangles indicate a significant difference between the null model prediction and the actual value, while circles indicate no significant difference. Bottom panels show the unpaired mean difference in litter decomposition rate between each treatment and the control group, with circles/triangles representing the effect size (mean difference) and its 95% confidence interval; triangles indicate a significant difference from the control (95% CI excluding zero), while circles indicate a non-significant difference.

The combined six-factor treatment produced the strongest suppression of the two biological processes, soil respiration and litter decomposition rate, across all four landscape elements. Respiration declined to only 82.40 ± 29.70–198.01 ± 48.67 ppm CO₂, while litter decomposition rate declined to 10.73 ± 5.38–21.81 ± 5.01%. This is consistent with the finding of Rillig, et al. ^1^, who suggested that increasing numbers of co-occurring GCFs produce increasingly directional changes in ecosystem functioning that cannot be predicted from single-factor responses alone.

Our results demonstrate that the four landscape elements, despite being co-located within the same region and exposed to identical climatic and experimental conditions, responded very differently to the same global change factors. This finding supports Hypothesis 1, that soil origin determines not only the baseline functional state of soils but also their sensitivity to multiple global change factors.

### 3.2. Non-additive interactions among multiple global change factors

Null model analyses revealed that some responses of soil functions to the combined six-factor treatment could not be consistently predicted by any null model derived from the single factor treatments, while some response variables in some soils fit additive, multiplicative, or dominative models.

Among the five response variables, electrical conductivity exhibited the highest predictability. In all four soils, the observed response of the six-factor treatment was consistent with the additive null model, indicating that the effects of individual stressors accumulated approximately linearly (**Figure S3**). This pattern is not unexpected because electrical conductivity is primarily a physicochemical property determined by the concentration of dissolved ions^36^. Therefore, it is more likely to follow additive expectations. In contrast, several other variables deviated from null model predictions. For water-stable aggregates, grassland and oak forest soils closely matched the model predictions (**Figure 2**). However, farmland and pine forest soils resulted in significantly higher values than predicted by any of the three null models. This suggests that the negative effects of individual stressors did not accumulate linearly in farmland and pine forest soils. Instead, antagonistic interactions among stressors may have partially buffered aggregate breakdown^11^. Such antagonistic responses may reflect the resistance and resilience of soil microbial communities, which can buffer ecosystem functioning against multiple disturbances^37^. A similar pattern was observed for soil pH, where neither oak forest nor pine forest soils followed any of the null model predictions (**Figure S2**). The observed pH remained closer to the control than expected. These deviations from the model predictions indicate that soil physicochemical buffering and interactions among multiple stressors can substantially alter the overall response under combined environmental change.

Soil respiration and litter decomposition rate showed non-additive responses. Only farmland soil followed the additive prediction for respiration, while the remaining soils were better described by multiplicative or dominative models (**Figure 3**). Oak forest and pine forest soils followed different null models for litter decomposition rate, while farmland and grassland soils exhibited decomposition rates that were lower than predicted by all three models (**Figure 4**). These results indicate that the inhibitory effects of multiple global change factors on decomposition exceeded those expected from the individual stressors alone. Such responses are consistent with synergistic interactions among stressors acting on soil microbial communities. Simultaneous exposure to warming, drought, salinity, antibiotics, microplastics, and nitrogen deposition may reduce soil biodiversity and functioning^38^. These stresses are likely to exceed the tolerance of microbial decomposers, producing disproportionately strong reductions in respiration and litter decomposition.

These findings support Hypothesis 2 and demonstrate that the effects of multiple global change factors cannot be universally predicted from the responses to individual stressors alone. These findings are consistent with experimental and meta-analytical evidence showing that ecosystem responses to multiple global change drivers are frequently non-additive, with both synergistic and antagonistic interactions causing responses to deviate from predictions based on individual stressors^39–41^. In soil ecosystems, such non-additive responses are due to multiple stressors simultaneously altering soil physicochemical conditions, resource availability, and microbial activity^42,43^, thereby generating interactions that cannot be predicted from single-factor experiments alone.

### 3.3. Multiple stressors amplified landscape-element specific response patterns

We performed principal component analysis of Hedges’ g effect sizes to characterize and compare the overall response patterns of the four soils to single and combined GCF treatments. **Figure 5a** showed clear separation among soils, especially between forest (oak and pine forests) and non-forest (farmland and grassland) soils. PERMANOVA analysis confirmed that soil explained substantially variance in the multivariate response (*p* = 0.006). Separation of soils was weak under single-stressor conditions, while it became more pronounced under the combined treatment. **Figure 5b** showed clear separation among treatments (*p* = 0.001), with multifactor treatment clearly separated from all single factor treatment. This suggests that landscape element-dependent responses were amplified under multi-stressor conditions.

**Fig. 5.**
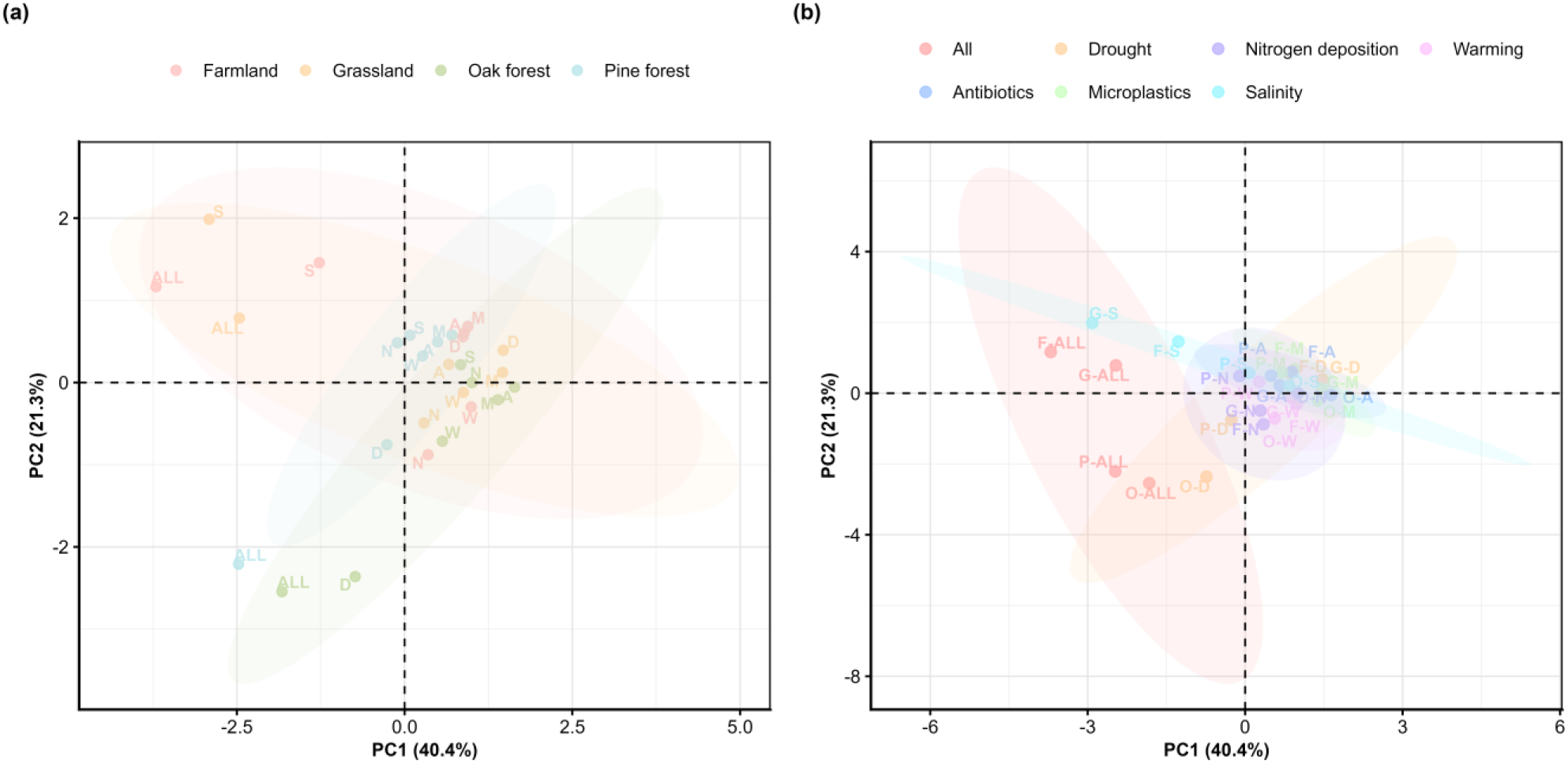
Principal component analysis of Hedges’ *g* values across (a) soils from four landscape elements and (b) global change factor treatments. Each point represents the multivariate effect size (Hedges’ *g*) of a single treatment within soil from one landscape element. (a) Points of each landscape elements cluster together and form a ellipse (*p* = 0.006). (b) Points of each treatment cluster together and form an ellipse (*p* = 0.001). Points are colored according to (a) soil or (b) treatment, and shaded ellipses indicate the 95% confidence regions for each (a) soil or (b) treatment group. PC1 and PC2 explain 40.4% and 21.3% of the total variance, respectively. A – antibiotics, All – six-factor treatment, D – drought, M – microplastics, N – nitrogen deposition, W – warming.

Mantel tests were conducted to examine associations between the effect size (Hedges’ *g*) of the five response variables and 19 soil physicochemical properties (**Figure S4**). Most of the Mantel tests did not detect significant associations between response-variable effect sizes and soil physicochemical properties. The only significant association was between the water-stable aggregates effect size and soil electrical conductivity (Mantel’s *r* = 0.95, *p* = 0.042). No other response variable–property combination was statistically significant. Thus, even though there is a significant separation among landscape elements in the multivariate analysis, no consistent relationship between physicochemical differences and differences in response patterns was found. This suggests that the individual measured properties may not capture the full set of factors underlying the context dependency observed among the four landscape elements.

One possible explanation is that landscape-specific responses are partly shaped by other factors that were not directly measured in this study. Forest and non-forest soils differ not only in their physicochemical properties but also in litter inputs, disturbances, land management history, and microbial community composition^44^. These factors collectively shape how soils respond to environmental disturbance. Previous studies have shown that forest and agricultural soils often differ substantially in microbial community composition, which can strongly influence ecosystem responses to environmental change^45^. Furthermore, soil properties and microbial communities interact through bidirectional feedbacks, with soil conditions shaping microbial communities and microorganisms in turn modifying soil physical, hydrological, and chemical properties^46^. Land-use history has also been shown to have a stronger influence on soil microbial community composition than aboveground vegetation and soil physicochemical properties^47^. Therefore, direct measurements of microbial community composition, functional traits, and functional redundancy would be needed to identify the mechanisms underlying the landscape-specific response patterns observed in this study.

## 4. Conclusion and ecological implications

We found that soils originating from different landscape elements in our case study area exhibit distinct sensitivities and response patterns when exposed to the same global change factors. This study provides direct experimental comparison of soils from contrasting landscape elements under multiple global change factors. We further demonstrate non-additive interactions among multiple global change factors, and that the magnitude and direction of these deviations from null model predictions are themselves context dependent. Furthermore, landscape-element specific response patterns were amplified under multiple stressors.

Our findings show that neighboring landscape elements can exhibit contrasting capacities to resist global change despite sharing the same regional environmental conditions. Landscape heterogeneity therefore represent a mosaic of stability and resilience. These findings highlight the value of integrating co-located landscape elements into multifactor global change experiments. Future experiments should therefore combine multifactorial treatments with comparisons among co-located landscape elements to better capture the complexity of ecosystem responses under realistic future scenarios. Finally, our results also have important implications for ecosystem policy and management, as approaches based on a single representative soil or land-use type may overlook substantial within-landscape variation in ecosystem responses. Soil responses should therefore be interpreted within the broader ecological and landscape context in which they occur^48^. Future global change assessments and management strategies should better incorporate landscape-level heterogeneity and consider the specific stability and resilience of individual landscape elements^49^. Similarly, management strategies aimed at improving ecosystem resilience under global change should consider landscape elements individually.

## Acknowledgments

We thank Dr. Marcus Wicke, Zwillenberg-Tietz Foundation, Linde Research Station (Märkisch Luch, State of Brandenburg, Germany) for providing the soil and sampling site information. We thank Anika Lehmann, Anja Wulf, Sabine Buchert and Yuanze Sun for helping with the experiment. T.H. acknowledges the China Scholarship Council for a scholarship (202406260044).

## Data availability

All data generated during this study are available in the Figshare repository, https://doi.org/10.6084/m9.figshare.33395680.

## Code availability

The underlying code for this study is available in Figshare and can be accessed via this link https://doi.org/10.6084/m9.figshare.33399484.

## Authors contribution

T.H.: conceptualization, methodology, investigation, formal analysis, data curation, visualization, writing - original draft. M.H.B.: validation, writing - review & editing. M.C.R.: conceptualization, resources, funding acquisition, writing - review & editing.

