## supplementary material for "Soils from different landscape elements diverge in response to multiple global change factors"

**This file includes:**

Figures S1 to S4

Tables S1 to S2

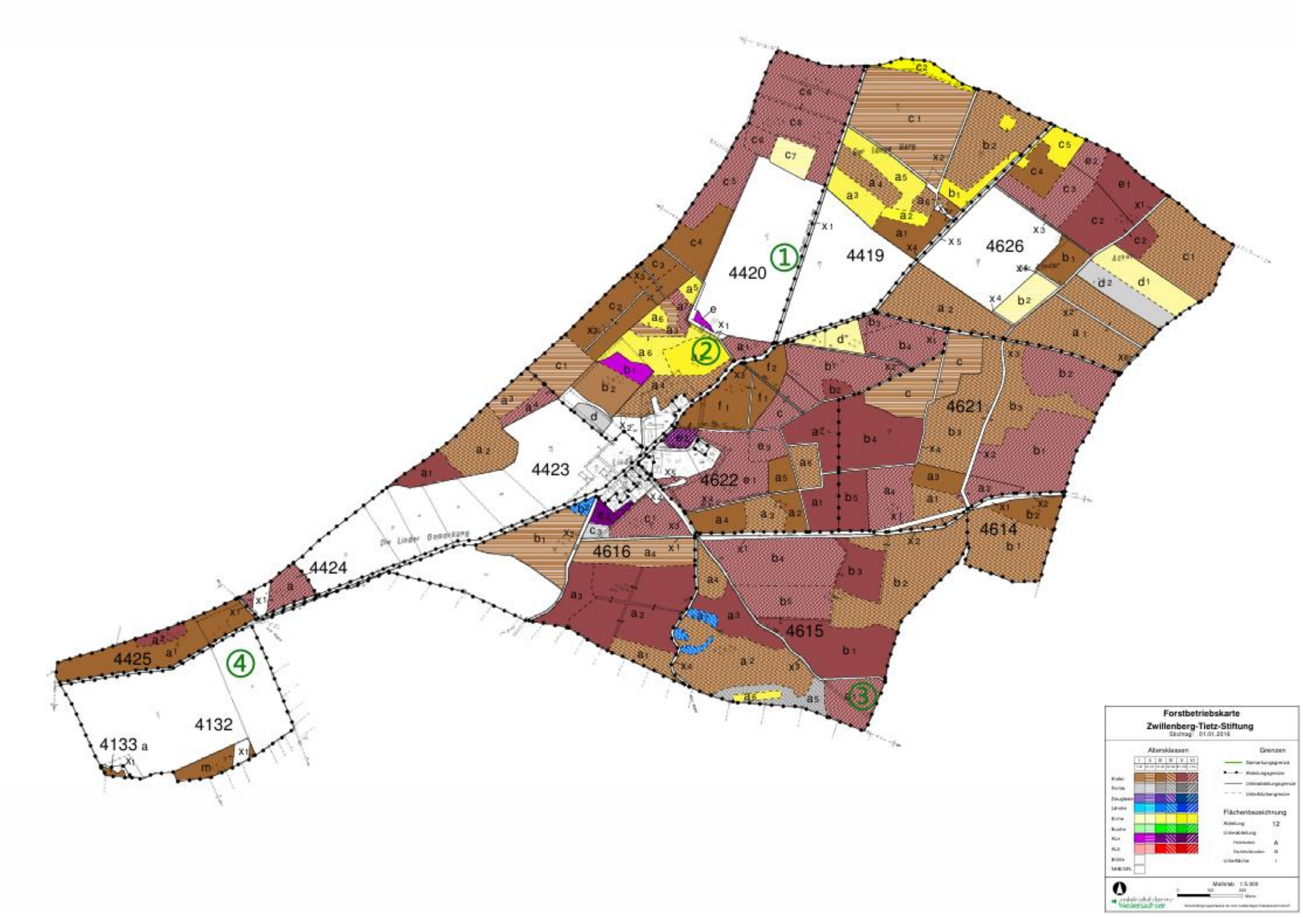


**Fig. S1 Sampling locations.** 1. farmland; 2. oak forest; 3. pine forest; 4. grassland. Vegetation map provided by Zwillenberg–Tietz Foundation.

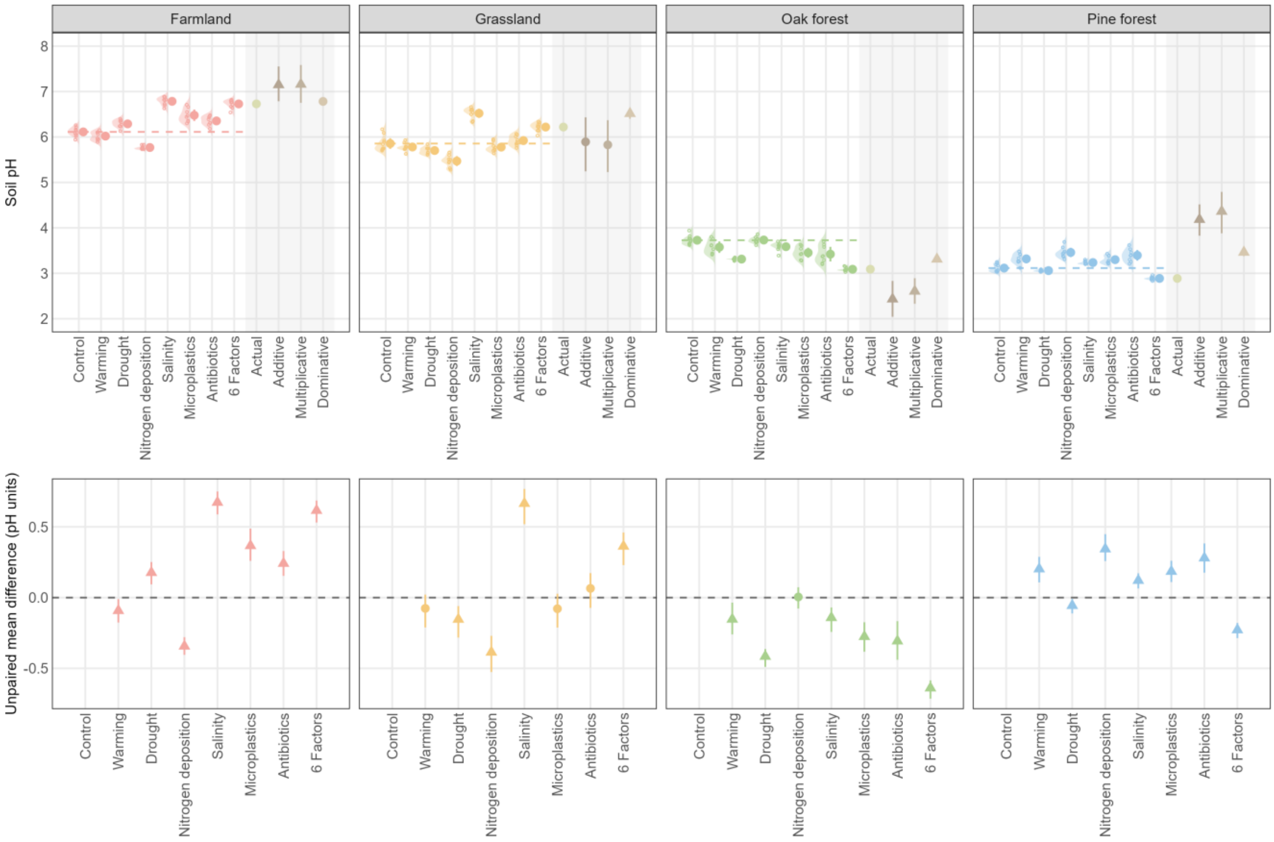


**Fig. S2. Responses of soil pH to individual and combined global change factor treatments across four landscape units.** Top panels show the raw data distributions (half-violin plots and jittered points) together with treatment means ± 95% confidence intervals. The dashed horizontal line indicates the mean pH of the control group. The grey-shaded area shows the observed ("Actual") pH under the combined six-factor treatment and the values predicted by three null models (Additive, Multiplicative, and Dominative); triangles indicate a significant difference between the null model prediction and the actual value, while circles indicate no significant difference. Bottom panels show the unpaired mean difference in pH between each treatment and the control group, with circles/triangles representing the effect size (mean difference) and its 95% confidence interval; triangles indicate a significant difference from the control (95% CI excluding zero), while circles indicate a non-significant difference.

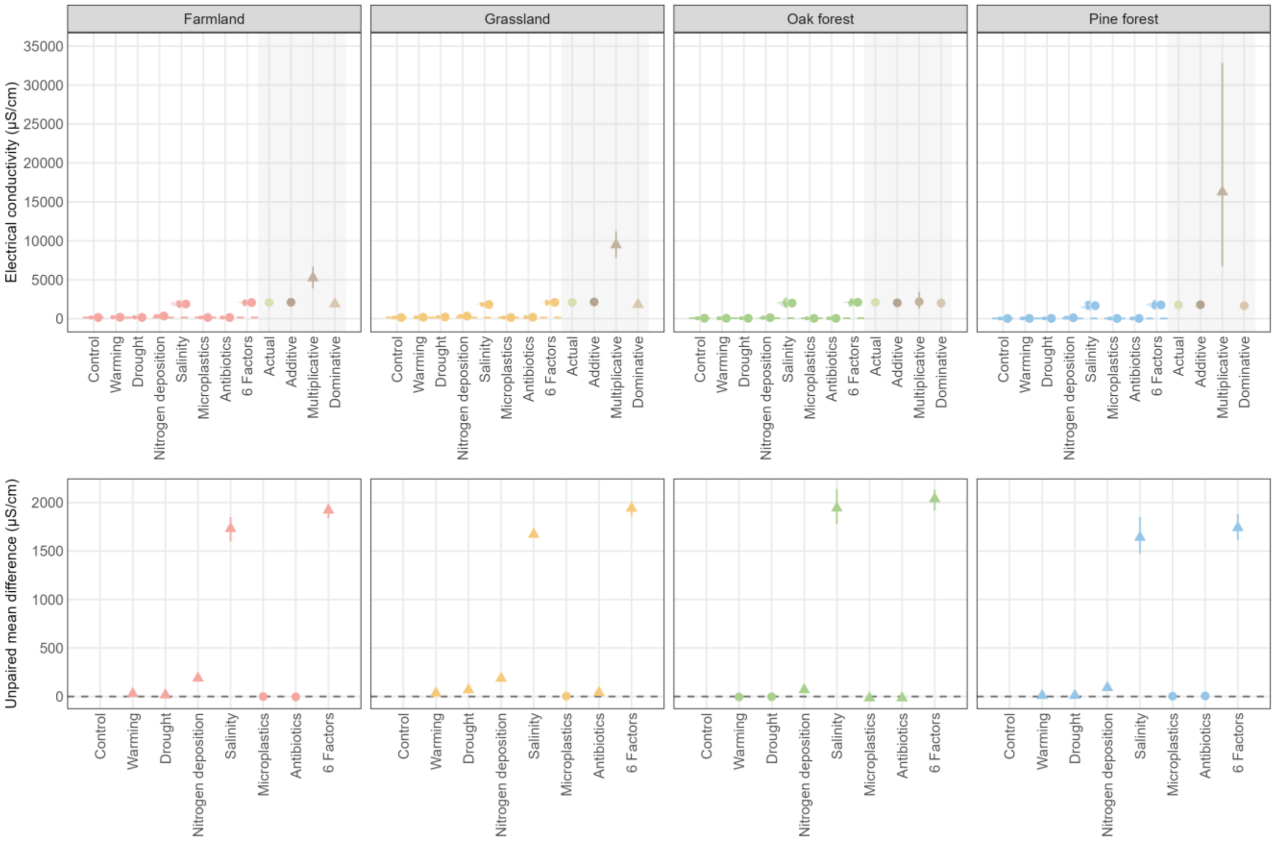


**Fig. S3. Responses of soil electrical conductivity to individual and combined global change factor treatments across four landscape units.** Top panels show the raw data distributions (half-violin plots and jittered points) together with treatment means ± 95% confidence intervals. The dashed horizontal line indicates the mean EC of the control group. The grey-shaded area shows the observed ("Actual") EC under the combined six-factor treatment and the values predicted by three null models (Additive, Multiplicative, and Dominative); triangles indicate a significant difference between the null model prediction and the actual value, while circles indicate no significant difference. Bottom panels show the unpaired mean difference in EC between each treatment and the control group, with circles/triangles representing the effect size (mean difference) and its 95% confidence interval; triangles indicate a significant difference from the control (95% CI excluding zero), while circles indicate a non-significant difference.

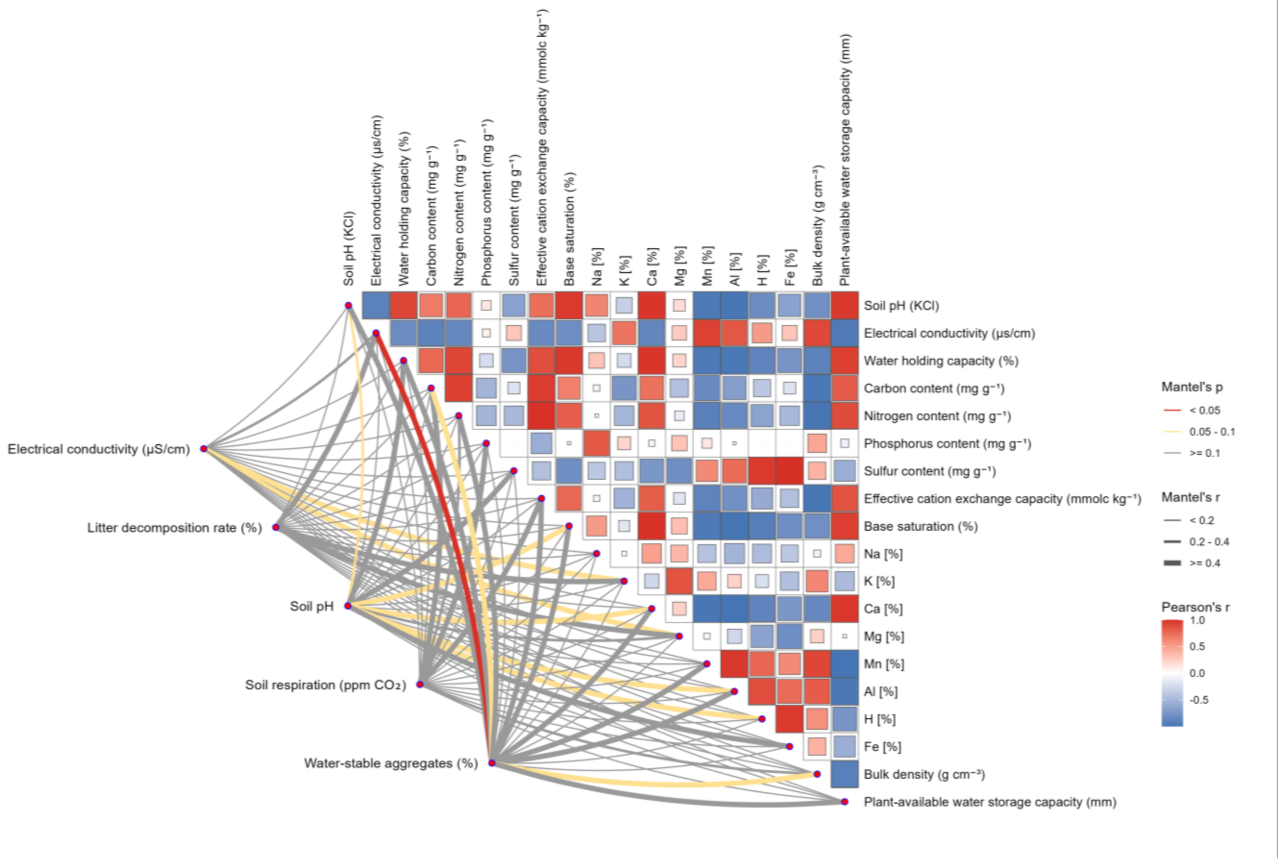


**Fig. S4. Mantel test correlations between soil physicochemical properties and response variable effect sizes**. Upper triangle: Pearson correlation matrix among 19 soil properties across four soil types. Lines connect each response variable to soil properties, with line color and width representing Mantel's p and r values, respectively.

**Table S1** Physicochemical properties of each soil sample. Data shown in italics were obtained from Pampe and Pampe [1]master thesis.

| Property | farmland | oak forest | pine forest | grassland |
|---|---|---|---|---|
| Soil pH (KCl) | 6.44 | 3.56 | 3.14 | 5.96 |
| Electrical conductivity (µs/cm) | 42.15 | 58.10 | 51.84 | 40.44 |
| Water holding capacity (%) | 28.48 | 18.96 | 12.66 | 34.13 |
| Carbon content (mg $g^{-1}$) | *5.5* | *2.7* | *4.7* | *8.7* |
| Nitrogen content (mg $g^{-1}$) | *0.5* | *0.3* | *0.3* | *0.8* |
| Phosphorus content (mg $g^{-1}$) | *0.3* | *0.2* | *0.2* | *0.1* |
| Sulfur content (mg $g^{-1}$) | *0.1* | *0.1* | *0.3* | *0.1* |
| Effective cation exchange capacity (mmolc $kg^{-1}$) | *26.2* | *10.8* | *12.9* | *54.0* |
| Base saturation (%) | *97.8* | *47.0* | *15.9* | *99.2* |
| Na [%] | *1.5* | *0.7* | *0.6* | *0.6* |
| K [%] | *3.7* | *7.7* | *2.3* | *2.3* |
| Ca [%] | *87.8* | *32.5* | *11.3* | *93.0* |
| Mg [%] | *4.8* | *6.1* | *1.8* | *3.4* |
| Mn [%] | *1.1* | *4.0* | *4.2* | *0.5* |
| Al [%] | *1.2* | *45.7* | *64.4* | *0.2* |
| H [%] | *0.0* | *3.3* | *13.5* | *0.0* |
| Fe [%] | *0.0* | *0.0* | *2.0* | *0.0* |
| Bulk density (g $cm^{-3}$) | *1.4* | *1.5* | *1.47* | *1.3* |
| Plant-available water storage capacity (mm) | *70.4* | *20.8* | *21.0* | *77.0* |

**Table S2** Type II ANOVA results for individual soil response variables.

| term | variable | eta_sq | Significance |
|---|---|---|---|
| soil | delta_EC | 0.08380574 | *** |
| treatment | delta_EC | 0.9851688 | *** |
| soil:treatment | delta_EC | 0.25326529 | *** |
| Residuals | delta_EC | | |
| soil | delta_LDR | 0.46080619 | *** |
| treatment | delta_LDR | 0.85604217 | *** |
| soil:treatment | delta_LDR | 0.49615356 | *** |
| Residuals | delta_LDR | | |
| soil | delta_pH | 0.8083356 | *** |
| treatment | delta_pH | 0.66164238 | *** |
| soil:treatment | delta_pH | 0.85939287 | *** |
| Residuals | delta_pH | | |
| soil | delta_CO2 | 0.49039265 | *** |
| treatment | delta_CO2 | 0.81989622 | *** |
| soil:treatment | delta_CO2 | 0.73150244 | *** |
